# Left Area PF as a Neural Marker of Technical Reasoning

**DOI:** 10.1101/2025.06.04.657853

**Authors:** Giovanni Federico, Ciro Rosario Ilardi, Paola Marangolo, Chloé Bryche, Maximilien Metaireau, Alexandre Bluet, Mathieu Lesourd, Yves Rossetti, François Osiurak

## Abstract

Humans possess a unique capacity for technical reasoning—the ability to infer and manipulate the causal structure of the physical world. Although this faculty is central to technological innovation, its neural substrates remain incompletely understood. Here, we show that grey matter volume in the left cytoarchitectonic area PF, within the supramarginal gyrus of the inferior parietal lobule, *selectively* predicts and *accurately* classifies technical reasoning performance in healthy adults (N = 75; 54 females; mean age = 20.92 ± 3.28 years). This association persists independently of demographic factors, personality traits, and total brain volume. By contrast, grey matter volume in right prefrontal regions, examined as control areas, correlates with broader cognitive functions—namely, fluid intelligence and abstract reasoning, but not with technical reasoning. These findings support the hypothesis that the left area PF is a domain-specific substrate for technical reasoning. Situated within a parietal territory that is evolutionarily expanded in humans, this region may constitute a neural signature of the human capacity to understand and reshape the material world.

## INTRODUCTION

Human beings possess a singular capacity to reshape the physical world. From crafting the earliest stone tools to designing skyscrapers and interplanetary probes, the evolutionary success of our species has hinged on a distinctive cognitive faculty: the ability to understand, manipulate, and innovate within the physical world.

This faculty, broadly conceived as *technical reasoning*, enables individuals to infer mechanical principles and devise functional solutions to real-world problems. In many ways, it forms the cognitive scaffolding upon which technological civilisation is built (*1, 2*).

Identifying the neural architecture that underpins this form of causal reasoning is critical to understanding the biological basis of a defining human trait. Comparative and evolutionary neuroscience have pointed to reorganisation within the parietal cortex, particularly the left inferior parietal lobule, as a key substrate for physical reasoning and material innovation (*3*). Within this territory, the left cytoarchitectonic area PF (Parietal F; **Figure 1**), located in the anterior supramarginal gyrus (*4*), has been consistently associated with the conceptualisation, planning, and manipulation of human-made artefacts (*5*–*7*).

**Figure 1.**
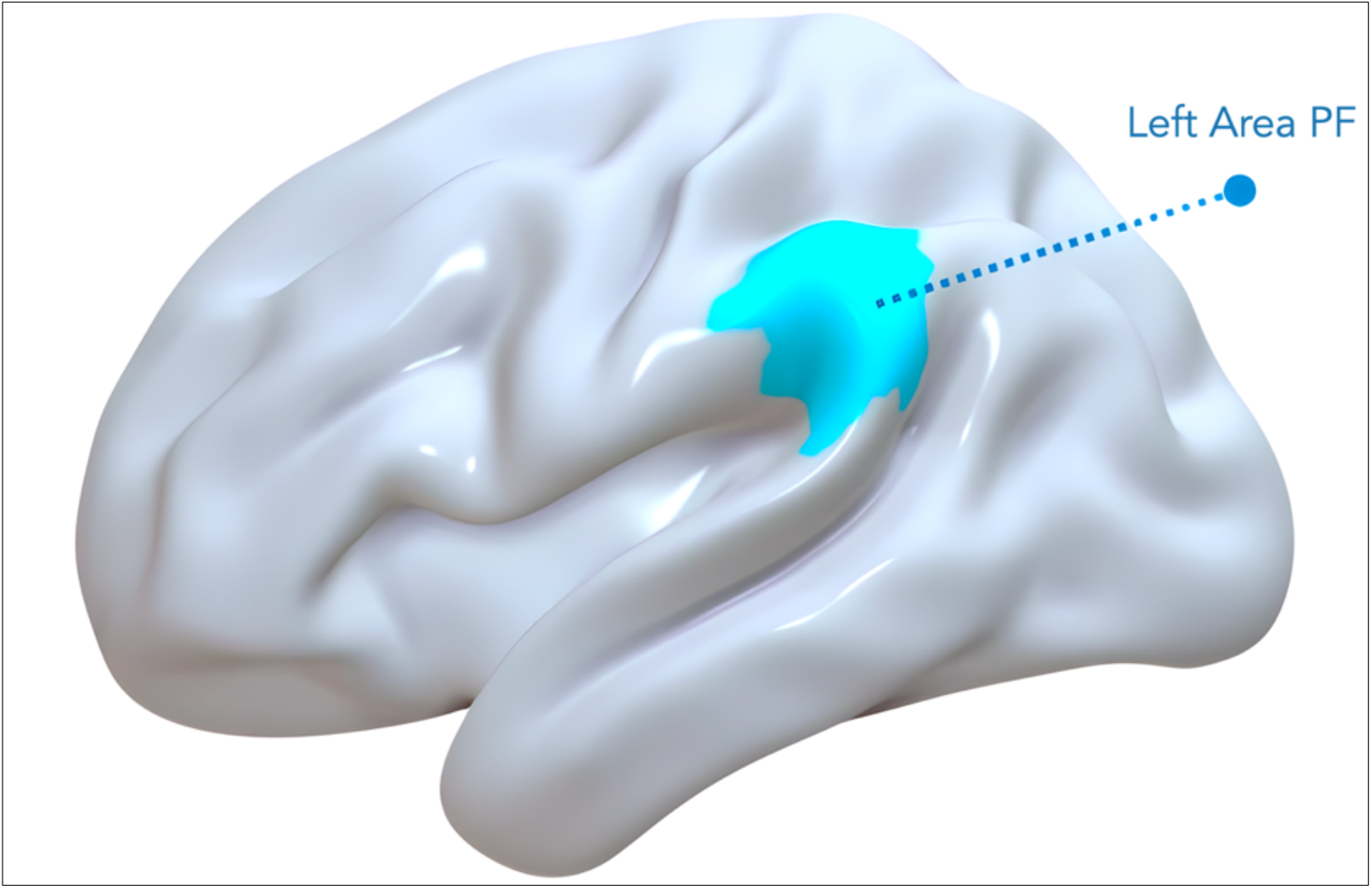
The Left Area PF. The area PF, located in the anterior supramarginal gyrus of the left Inferior Parietal Lobe and defined according to (*36*), is depicted in light blue.

This region has been robustly engaged across a wide array of studies examining physical problem solving and mechanical cognition (*5, 8*–*15*). Preliminary findings further suggest that individual differences in the morphology of this brain area may account for variability in technical reasoning performance (*16*). Clinical evidence supports this view: lesions to the left area PF result in *apraxia of tool use*, an acquired inability to execute meaningful actions with familiar tools, underscoring its central role in physical understanding (*17*–*22*). Yet, a key question remains unresolved: does the engagement of the left area PF reflect a *specialised* neural mechanism evolved to support technical reasoning, or does it simply mirror the recruitment of broader cognitive faculties, such as fluid intelligence or general problem-solving ability?

To disentangle these possibilities, we investigated whether anatomical variation in the left area PF *specifically* predicts technical reasoning performance, as distinct from broader intellectual abilities. Using structural magnetic resonance imaging, we measured grey matter volume, a commonly used indicator of regional neural structure and its relation to cognitive function (*23*–*25*), in a sample of healthy adults. We then examined the relationship between neuroanatomical data and behavioural performance on a comprehensive neuropsychological battery designed to examine technical reasoning, fluid intelligence, and problem-solving skills.

Our analyses controlled for potential confounds including age, sex, handedness, total intracranial volume, and personality traits. To assess anatomical specificity, we conducted comparative analyses in right inferior frontal regions—a prefrontal territory traditionally linked to general reasoning and executive processes, but not typically associated with technical reasoning (*26*–*28*). Lastly, we tested whether grey matter volume in the left area PF could reliably distinguish individuals with high versus low technical reasoning ability, offering a potential neural marker of this cognitive trait.

## METHODS

The methods and procedures employed in this study, particularly those related to MRI imaging, the analytical approach, technical reasoning tasks, and associated metrics, follow those outlined in (*16*).

### Participants

Seventy-five right-handed participants (females = 54; mean age = 20.92 ± 3.28 years) were recruited from Lyon University, France. Recruitment criteria included: (i) no current or history of alcohol or drug abuse; (ii) no current or history of major psychiatric disorders; (iii) no history of brain injury, stroke, or other primary clinical conditions; and (iv) no current or past use of psychoactive medications. These criteria were assessed through a clinical interview conducted by an experienced medical doctor. All participants provided written informed consent before participation in the study.

### Procedure

The study was carried out at the Laboratory for the Study of Cognitive Mechanisms (EA 3082) at the University of Lyon and the Lyon Neuroimaging Department (CERMEP) in Lyon, France. All procedures complied with the ethical guidelines of the 1964 Declaration of Helsinki. Approval for the study was obtained from the French Ethics Committees (approval number: IRB: 2020-A02115–34). Participants were recruited randomly via social media advertisements. Before the scheduled MRI session, participants signed an informed consent form and underwent a medical evaluation to confirm their suitability for the MRI procedure. They then completed the neuropsychological battery at the Laboratory for the Study of Cognitive Mechanisms. The MRI session was conducted at CERMEP.

### Neuropsychological tests

A comprehensive neuropsychological battery was administered to evaluate distinct problem-solving and reasoning abilities. Handedness was measured using the *Edinburgh Handedness Inventory* to account for potential lateralisation effects on cognitive performance (*29*). Personality traits were assessed using the *Big Five Questionnaire* (BFQ; Extraversion, Agreeableness, Conscientiousness, Neuroticism, Openness; John et al., 2012). Non-verbal reasoning and abstract thinking were evaluated with *Raven’s Progressive Matrices*, a widely employed test of fluid intelligence (*31*). Technical reasoning was assessed using two subtests from the *NV7 psycho-technical battery* (*32*): the Physical Properties subtest (24 items, assessing understanding of physical properties) and the Visuospatial Imagery subtest (38 items, requiring identification of three-dimensional shapes from two-dimensional patterns). These scores were normalised and used to derive a *Technical Reasoning Performance Index* (TRPI), which was calculated according to the following equation (*16*):

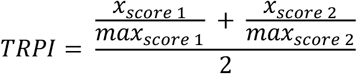

where *x*_*score* 1_ and *x*_*score* 2_ were the participants’ scores (*i*.*e*., physical-understanding and visuospatial-imagery score, respectively); *max*_*score* 1_ and *max*_*score* 2_ were the maximum obtainable scores at the two NV7 subtests (*i*.*e*., 24 points for the physical-understanding skills and 38 points for the visuospatial-imagery abilities). The TRPI varied between 0 and 1, where 0 was the lower technical-reasoning performers and 1 the higher ones. Finally, participants completed the *D2000 Domino Test*, a psychometric tool that gauges general (*g*) and fluid intelligence through problem-solving tasks involving domino-like figures (*33*). Each item presents a sequence of domino pieces, requiring identifying the underlying logical rule to select the piece that completes the pattern. The D2000 assesses multiple cognitive processes, including numerical reasoning (arithmetic operations and numerical relationships among domino pips), spatial reasoning (manipulating spatial arrangements of domino pieces), and abstract reasoning (isolating logical patterns beyond simple spatial or numerical cues).

### MRI scanning and brain morphometry analysis

Three-dimensional (3D) T1-weighted (T1w) images were acquired using a 3-Tesla Siemens Prisma scanner with a 64-channel head coil. The imaging parameters were as follows: repetition time (TR) = 3000 ms, echo time (TE) = 2.93 ms, flip angle = 8°, voxel size = 0.8 × 0.8 × 0.8 mm, matrix = 280 × 320, and field of view = 224 × 256. Data were collected in DICOM format and converted to NIfTI format using the *dcm2niix* software. All images underwent visual inspection by an experienced neuropsychologist (G.F.) to identify artefacts. We employed *FastSurfer*, a deep-learning-based neuroimaging pipeline, to process T1-weighted structural MRI data (*34*). *FastSurfer* offers a rapid and accurate alternative to traditional neuroimaging tools, providing outputs compatible with *FreeSurfer* (*35*). *FastSurfer*’s segmentation and surface reconstruction pipelines were employed to obtain each subject’s cortical and subcortical segmentations and cortical surface models. Subsequently, the *Human Connectome Project Multi-Modal Parcellation* (HCP-MMP) atlas was applied to each subject’s cortical surface (*36*). This involved mapping the atlas annotations from the *fsaverage* template to the individual subject’s anatomy for both left and right hemispheres using the *mri_surf2surf* utility. Cortical statistics for each region defined by the HCP-MMP atlas were then computed using the *mris_anatomical_stats* tool, providing region-specific morphometric data for further analysis. The structural MRI analysis employed in this study aligns with the one detailed in (*16*), with the primary modification being the utilisation of *FastSurfer* in place of *FreeSurfer* for segmentation and surface reconstruction.

### Statistical Analyses

One-tailed, ninth-order *Pearson’s partial correlations* 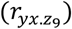 were conducted to examine the associations between neuropsychological scores and grey matter volumes (GMV) in the right inferior frontal junction *area a* (IFJa), right frontal operculum *area 5* (FOP5), and left *area PF*, while controlling for nine covariates. These included sex, age, Edinburgh Handedness Inventory score, estimated total intracranial volume, and the five subscales of the Big Five Questionnaire (BFQ). The primary endpoint was to quantify the partial correlation between the TRPI score and PF GMV. Additionally, the associations between IFJa GMV and Raven’s Progressive Matrices and FOP5 GMV and D2000 were examined as a control analysis. For each correlation coefficient, the effect size was interpreted according to Cohen’s conventions: small 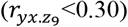, moderate 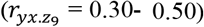, and large 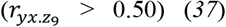. Bonferroni’s correction was used to mitigate the inflation of Type I error in multiple comparisons by appropriately adjusting the nominal alpha level (*α*_*adj*_ = 0.008).

Subsequently, we implemented a *forward regression analysis* to understand better the individual and combined contributions of candidate predictors in explaining the variance of PF GMV. To account for potential nonlinear relationships, each putative predictor was first evaluated for the best-fitting functional form, testing multiple regression models, such as linear (*Y* = *β*_0_ + *β*_1_*x* + ε), inverse 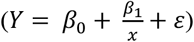, quadratic 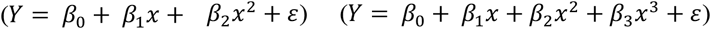, and logarithmic (*Y* = *β*_0_ + *β*_1_*logx* + ε) models. Once the most appropriate transformation was identified for each predictor, the final regression model was constructed iteratively, with variables entered sequentially based on their contribution to the most significant increase in the explained variance (*R*^2^). Predictors were retained only if they significantly improved model fit, as determined by multiple goodness-of-fit indices. These included the Akaike Information Criterion (AIC) and Bayesian Information Criterion (BIC) (*38*). The Wilkinson-Dallal method was used to mitigate Type I error inflation (*39*). In line with the *F*-to-Enter Stopping Rule, critical *R*^2^ values were defined using α = 0.05, *k* candidate predictors, residual *df* = *N* − 1 − *k*, and *F*-to-enter = 4.00. Regression analysis was conducted upon verification of major Gauss-Markov’s generalised linear model assumptions, including the absence of severe multicollinearity, normally distributed residuals, homoscedasticity, and no significant autocorrelation (*40*). The multicollinearity assumption was tested using Variance Inflation Factors (VIFs) and tolerance values. VIF values below 5 and tolerance values above 0.1 confirmed the absence of multicollinearity (*41*). Normality of residuals was assessed by inspecting Q-Q plots, while homoscedasticity was visually confirmed by plotting standardised residuals against predicted values. Autocorrelation of residuals was tested using the Durbin-Watson statistics, with values close to 2 indicating no autocorrelation (*42*).

Having identified the optimal regression model, we subsequently examined the extent to which PF volumetry could discriminate between high and low performers on a technical reasoning task. To this end, we conducted a nonparametric receiver operating characteristic (ROC) curve analysis to assess discriminatory power. The state variable was constructed by assigning a value of 1 to individuals scoring below the 10th percentile on TPRI and 0 otherwise. Following established conventions, an area under the ROC curve (AUC) exceeding 0.70 indicated acceptable diagnostic accuracy. The optimal cutoff was determined using the Youden index (*J, sensitivity* + *specificity* − 1), and a comprehensive set of diagnostic accuracy metrics was computed (*43*).

Each quantitative variable was converted into *z* scores before running the analyses, as appropriate. Missing data (<3%) were handled using listwise deletion. Statistical analyses were performed using IBM SPSS Statistics v. 27 and R Studio v. 4.3.2.

## RESULTS

### Partial correlation analysis

The TRPI score demonstrated a moderate association with PF GMV 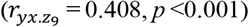, whereas no significant correlations were observed with IFJa 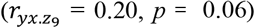 or FOP5 GMVs 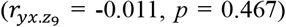. The Raven’s Progressive Matrices showed a moderate correlation with IFJa GMV 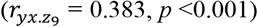 but no significant associations with PF 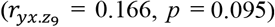 or FOP5 GMVs 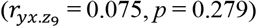. Finally, D2000 score exhibited a weak-to-moderate correlation with FOP5 GMV 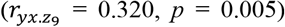, with no significant association with PF GMV 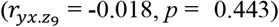. A weak correlation also emerged between D2000 score and IFJa GMV 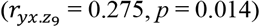; however, this correlation did not survive Bonferroni’s correction. The results of the partial correlation analyses are depicted in **Figure 2**.

**Figure 2.**
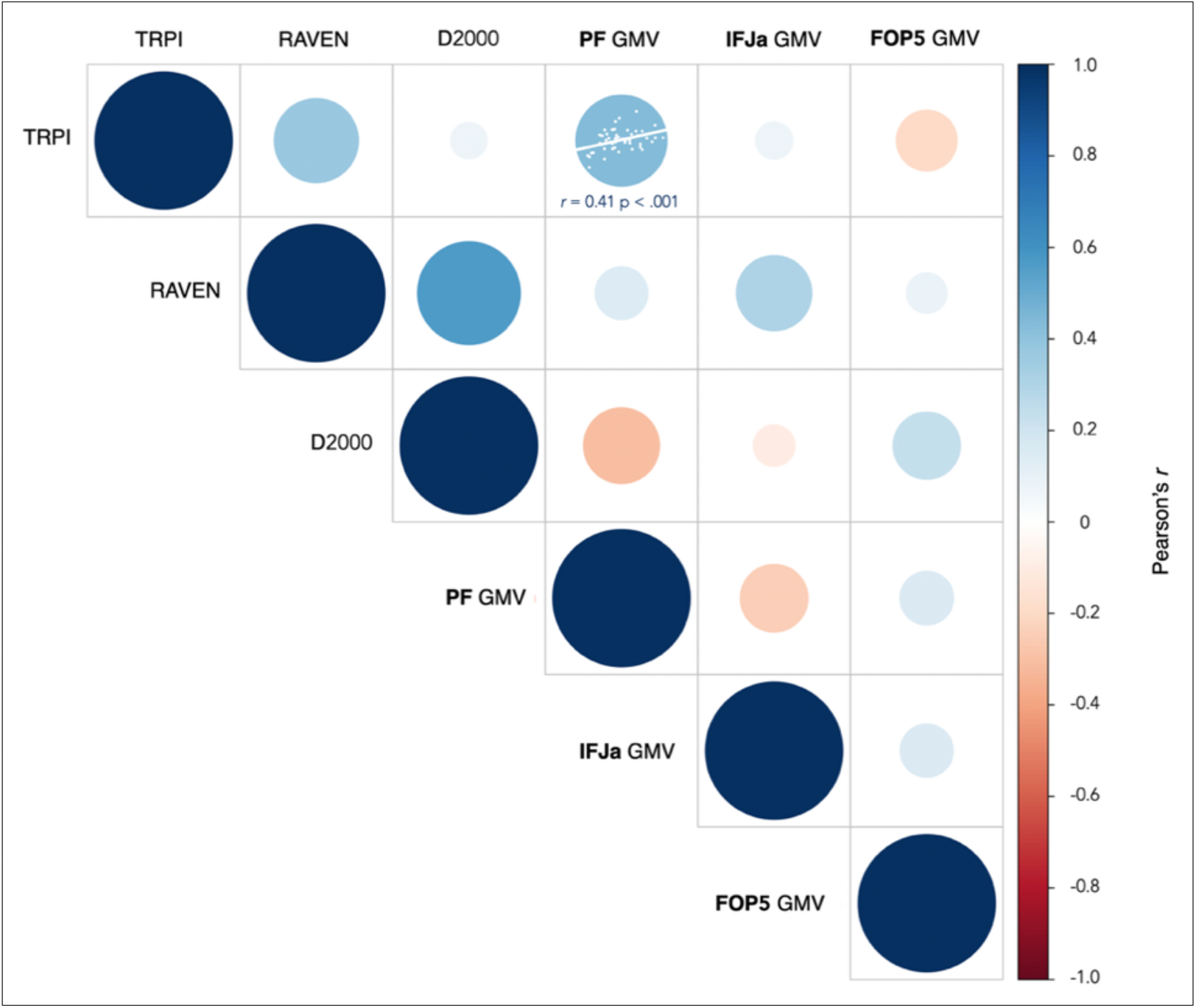
Pearson’s Partial Correlations. Pearson’s partial correlations are shown between scores on the Technical Reasoning Performance Index (TRPI), fluid intelligence (Raven’s Progressive Matrices; RAVEN), problem-solving abilities (D2000 test), and grey matter volume (GMV) in three brain regions: the left cytoarchitectonic area PF (PF), the right inferior frontal junction area (IFJa), and the right frontal operculum area 5 (FOP5). Each circle’s size and colour represent the correlation’s strength and direction (see scale bar).

### Multivariate hierarchical regression with log-linear, polynomial, and inverse terms

To capture the most accurate representation of the relationship between PF GMV and its predictors, each variable was tested against multiple models. TRPI score was best represented by a quadratic function, indicating a nonlinear, U-shaped relationship between TRPI and PF GMV (*R*^2^= 0.099, *F*_(2,72)_ = 3.974, *p* = 0.023; *β*_1_ = 0.250, *p* = 0.03; *β*_2_= −0.226, *p* = 0.048). Instead, an inverse function best described BFQ-Openness (*R*^2^= 0.099, *F*_(1,71)_ = 7.796, *p* = 0.007; *β* = 0.315, *p* = 0.007). All other tested variables—regardless of functional form—failed to reach statistical significance (*p* > 0.05) and were excluded from the final model. A forward regression model was built iteratively using a hierarchical approach. At stage 1, the TRPI score was entered to examine its primary contribution to the model. At stage 2, BFQ-Openness was included to assess whether personality traits explained additional variance beyond the cognitive predictor. TRPI was centred before polynomial expansion to mitigate multicollinearity. BFQ-Openness entered as an inverse transformation. As shown in **Table 1**, the final model included the TRPI score (linear term; see **Figure 3**) as the sole predictor, accounting for 6% of the variance in PF GMV (*F*_(1,71)_ = 4.283, *β* = 0.239, *p* = 0.042; AIC = 3.419, BIC = 0.996). The forward selection process did not retain the quadratic term and BFQ-Openness due to their negligible contribution to PF GMV variance (*p* > 0.05). According to Wilkinson and Dallal’s conservative test of significance, for α = 0.05, *k* = 3 predictors, residual *df* = 71, and the *F*-to-enter = 4.00, the critical *R*^2^ value ranged between 0.05 and 0.07. Therefore, our model can be regarded as statistically robust. Thus, a final regression equation was derived, serving as the mathematical distillation of our findings. Grounded in empirical evidence, this equation formalises the relationships that govern PF GMV variability:

**TABLE 1.** Results of Multivariate Hierarchical Regression.

| Predictor | $\beta$ | Student's t | p-value | Tolerance | VIF |
| --- | --- | --- | --- | --- | --- |
| Intercept |  | 0.072 | 0.943 |  |  |
| TRPI (linear) | 0.245 | 2.070 | 0.042 | 1.000 | 1.000 |
| TRPI (quadratic)<sup>e</sup> | -0.223 | -1.959 | 0.054 | 0.984 | 1.017 |
| BFQ-Openness (inverse)<sup>e</sup> | -0.002 | -0.021 | 0.983 | 0.994 | 1.006 |

Note. TRPI: Technical Reasoning Performance Index. BFQ: Big Five Questionnaire. VIF: Variance Inflation Factor. <sup>e</sup> Variable excluded.

**Figure 3.**
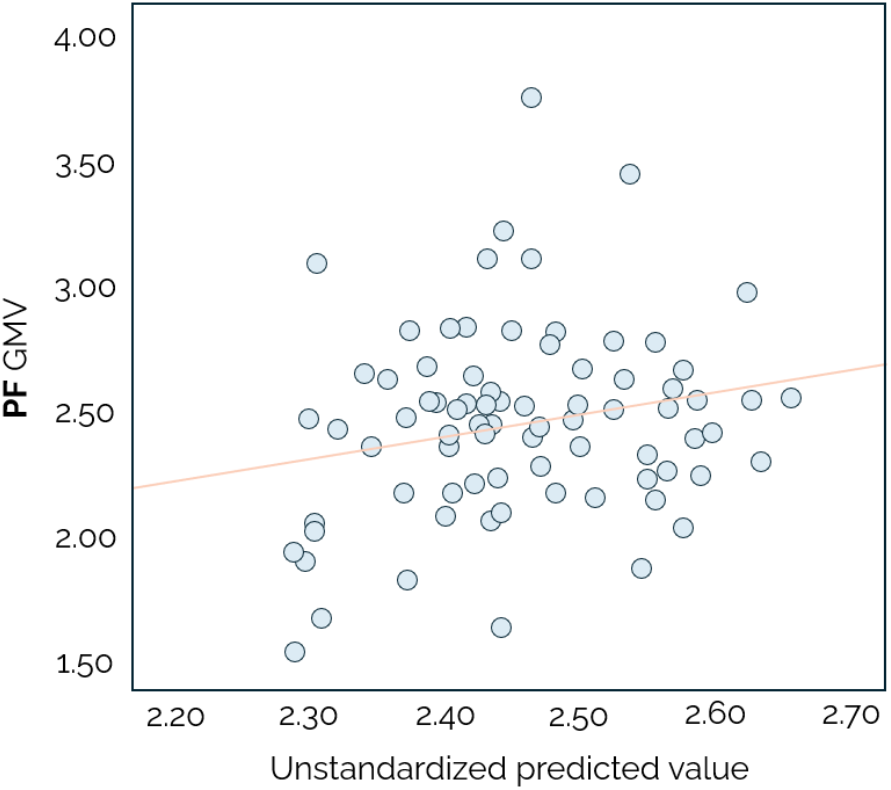
Association Between Grey Matter Volume in the Left Area PF and Technical Reasoning. Scatter plot illustrating the relationship between TPRI scores and grey matter volume in the left area PF (PF GMV). Data points reflect unstandardised predicted values derived from a multiple regression model.

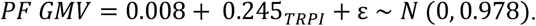

### Discriminatory power analysis

PF volumetry demonstrated adequate discriminatory power between participants who scored lower (*n* = 8) and higher (*n* = 67) on TPRI (AUC = 0.80 [95% CI 0.565-1.000], SE = 0.11, *p* <0.001; see **Figure 4a**). The optimal cutoff, determined using *J* for the best balance between sensitivity and specificity, was 2.054 (*J* = 0.705). This threshold value exhibited excellent sensitivity (Se = 0.750) and specificity (Sp = 0.955), with overall strong discriminatory power metrics, which are summarised in **Table 2**. Post-test probability is depicted as Fagan’s nomogram in **Figure 4b**.

**TABLE 2.** Discriminatory Performance of the Optimal Grey Matter Volume Cut Point in Left Area PF.

| Property | Value [95% CI] |
| --- | --- |
| Sensitivity | 0.750 [0.409-0.929] |
| Specificity | 0.955 [0.876-0.985] |
| False Positive Rate | 0.045 [0.015-0.124] |
| False Negative Rate | 0.250 [0.071-0.591] |
| Accuracy | 0.933 [0.853-0.971] |
| Positive Likelihood Ratio (LH+) | 16.750 [5.167-54.300] |
| Negative Likelihood Ratio (LH-) | 0.262 [0.079-0.870] |
| Post-Test Probability (Positive Predictive Value) | 0.667 [0.382-0.866] |
| Post-Test Probability (Negative Test) | 0.030 [0.009-0.094] |

**Figure 4.**
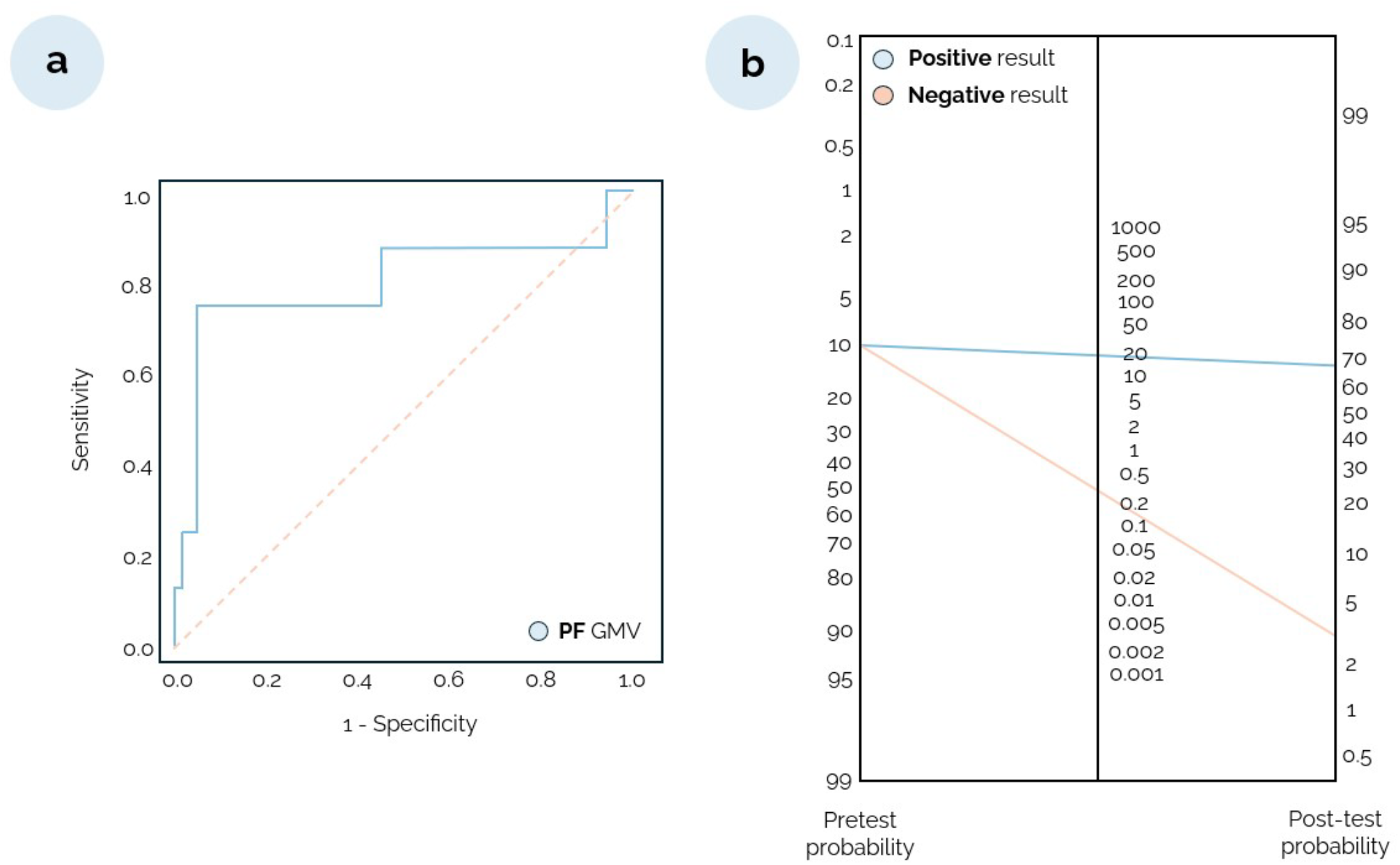
Discriminatory Power Analysis. (a) Receiver operating characteristic (ROC) curve showing the discriminative power of grey matter volume in the left area PF (PF GMV) for identifying individuals with low versus high technical reasoning ability, as measured by the TPRI (low: n = 8; high: n = 67). (b) Fagan’s nomogram illustrating post-test probabilities based on an optimal PF GMV threshold of Given a prevalence of ~10%, a positive test result yields a post-test probability of ~70% for correct classification as “atechnic performance.” In contrast, a negative test result yields a post-test probability of ~3%.

## DISCUSSION

The primary objective of this study was to investigate whether the morphological properties of the left area PF, located in the anterior supramarginal gyrus of the inferior parietal lobe (*4*), function not only as a *sensitive* (*16*) but also as a *specific* neural correlate of technical reasoning, that is, the human capacity to understand and manipulate the mechanical properties of the physical world (*2, 44*).

We found a selective association between grey matter volume in the left area PF and participants’ performance on behavioural tasks designed to capture technical reasoning abilities (*16, 32*). Crucially, this association did not extend to other cognitive domains, such as fluid intelligence and general problem-solving skills, which were assessed using two established neuropsychological instruments (*31, 33*). The specificity of this association was further substantiated by controlling for a range of potential confounds, including individual variability in brain morphology, socio-demographic factors, and personality traits (*30*).

By contrast, in line with prior evidence (*26*–*28*), brain regions within the right inferior frontal cortex were associated with performance on fluid intelligence and abstract reasoning/problem-solving tasks, but not with technical reasoning. Specifically, scores on Raven’s Progressive Matrices correlated with grey matter volume in the right inferior frontal junction *area a*, whereas general problem-solving performance (as indexed by the D2000 test) was associated with the frontal opercular *area 5* (*31, 33, 36*). Importantly, these right-frontal areas were included in the analyses as control sites, allowing us to delineate the domain specificity of the left area PF in supporting technical reasoning.

The cognitive specialisation of the left area PF is consistent with evidence indicating that the inferior parietal cortex is disproportionately expanded in *Homo sapiens* relative to non-human primates (*45*). This expansion co-occurs with the emergence of cumulative technological culture—the ability to create, refine, and transmit complex tools and mechanical systems across generations (*3*). While it is commonly held that such advances depend largely on imitation, teaching, and other forms of social learning (*46*–*50*), a complementary view proposes that these abilities are scaffolded by a domain-specific neurocognitive system for reasoning about physical causality, that is, technical reasoning (*2, 13, 44, 51*). Within this framework, the left area PF may represent the core neural substrate for this human technical faculty (*5, 10, 52*). Supporting this hypothesis, cytoarchitectonic studies indicate that structural homologues of the left area PF are largely absent in non-human species (*4*), underlining its potential evolutionary significance for technical cognition in *Homo sapiens*.

Converging neuroimaging evidence has implicated the left anterior supramarginal gyrus, including the area PF, in various tasks involving tool use and mechanical reasoning (*5, 8*–*15, 52*). Meta-analyses have shown that activity in this region increases when individuals focus on the mechanical structure of observed actions instead of their visuomotor aspects (*6*). Notably, this region is also activated during the passive observation of mechanical actions, suggesting that technical reasoning may be engaged vicariously (*53, 54*).

Clinical studies provide additional evidence for the unique role of the left area PF in technical reasoning. Research on *apraxia of tool use* in patients with left-hemisphere damage has revealed profound difficulties in manipulating familiar and novel tools to solve mechanical problems (*17*–*19, 55, 56*). These impairments, often characterised by an inability to infer tool-related physical properties or devise functional mechanical solutions, have been reliably associated with lesions in the left anterior supramarginal gyrus (*21, 22, 57, 58*).

Beyond establishing a robust structure–function relationship, our findings suggest that morphometric features of the left area PF may also exhibit *criterion validity*. Discriminatory power analyses revealed that grey matter volume in this region accurately distinguished individuals with high versus low technical reasoning abilities, with excellent sensitivity and specificity. This capacity for discrimination indicates that morphological indices in the left area PF may serve as a neuroanatomical marker of individual differences in technical reasoning. While further validation in independent cohorts is required, these results open promising avenues for translational applications in neurocognitive profiling and clinical neuroscience (*59*–*65*).

## CONCLUSION

This study identifies the left area PF as a specialised neural substrate for technical reasoning and demonstrates its predictive value for individual differences in technical reasoning. By revealing a selective association between the morphology of this region and technical reasoning, we propose that the left area PF enables humans to detect and manipulate causal regularities in the physical world. Such anatomical specialisation likely played a pivotal role in the emergence of cumulative technological culture throughout human evolution. Accordingly, the present findings lay the groundwork for future research at the intersection of cognitive neuroscience, evolutionary biology, and anthropology, aimed at further elucidating the origins and development of this human capacity.

## Author contributions

F.O. and G.F. developed the research question and designed the experiment. A.B., C.B., C.R.I., F.O., G.F., M.L., M.M., P.M., and Y.R. conducted the experiment. C.R.I., F.O. and G.F. analysed the data. G.F. wrote the manuscript with input from all the authors.

## Data availability

The data supporting the present study’s findings are available at https://osf.io/xq5sz.

